# Adaptation of *Acinetobacter baumannii* to colistin exposure: Laboratory mimicking of a clinical case

**DOI:** 10.1101/493221

**Authors:** Ezgi Oralkan, Siran Keske, Onder Ergonul, Fusun Can

## Abstract

Adaptation of *Acinetobacter baumannii* to colistin use on a disease course of a patient was described. Effects of colistin was mimicked in the laboratory conditions. The colistin resistant isolate was identified from the patient after 25 days of colistin treatment. In the laboratory, the expressions of *pmrCAB* were the highest at the generation which was corresponded to the duration of therapy. *A. baumannii* can develop a stable colistin-resistant phenotype after three weeks of colistin exposure.

---

The increase of colistin-resistance in *Acinetobacter baumannii* is a great concern in various regions of the world such as Asia, Europe, and North and South America (1). Exposure to colistin is considered as the most significant factor for emerging of colistin resistance (2), however, details of the *in vivo* response of *A. baumannii* to colistin exposure, such as the number of days for development of resistance was not described. The molecular studies about colistin resistance indicated two main mechanisms: modifications in lipid A structure and complete loss of LPS (3). By this study, we analyzed adaptation of *A. baumannii* to colistin use on a disease course of a patient. The cellular effects of colistin use was mimicked in laboratory conditions to understand the relationship between the clinical features and molecular resistance mechanisms.

A 45-year-old woman had a stab wound caused by penetration of the abdominal and thoracic walls by a sharp object. Two days after abdominal and thoracic surgery in another hospital, she was transferred to our intensive care unit (ICU). In 3^rd^ day of hospitalization, her body temperature was elevated, and her white blood cell count was 27,280 /µL, C-reactive protein was 356.9 mg/L and procalcitonin was 0.86 ng/mL. Carbapenem resistant *A. baumannii* (CarR-AB) was isolated from the cultures of sputum and abdominal drainage in 5^th^ day. Colistin 300 mg/day and meropenem 3 g/day were started, but clinical and laboratory response were poor. At the 15^th^ day of hospital stay, CarR-AB was isolated again from abdominal drainage. Meropenem was switched to tigecycline 100 mg/day and colistin was maintained. At the 29^th^ day, antibiotic treatment was stopped based on clinical and laboratory findings. At the 38^th^ day, colistin resistant *A. baumannii* was isolated from the abdominal drainage. *A. baumannii* was only susceptible to tigecycline and amikacin. Colistin 300 mg/day and tigecycline 100 mg/day were started, however colistin was switched to amikacin in 48^th^ day because of the generalized itching that does not response to anti-histaminic treatment. During the therapy with amikacin and tigecycline, she had generalized tonic-clonic seizures probably associated with amikacin in 50^th^ day. She had no fever, her white blood cell count was 12,420 /µL, C-reactive protein was 30.4 mg/L and procalcitonin was 0.24 ng/mL on the day of seizure and antimicrobial treatment was stopped. In the following days, she had no fever and she was discharged at the 72^nd^ day of hospitalization. At year 1 of follow-up, the patient was completely free of symptoms. At year 2 of follow-up, she was completely well, and follow-up laboratory tests were within normal limits.

Four colistin susceptible *A. baumannii* from sputum (K399, K411) and intra-abdominal fluid (IAF) (K408, K412) were isolated. After 25 days of colistin use, a colistin resistant *A. baumannii* (K409) was isolated from IAF. Clonal relatedness of the isolates was determined by the repetitive PCR (rep-PCR) based Diversilab system (Biomerux, France). The colistin minimum inhibitory concentration (MIC) of each isolate was determined by broth microdilution in accordance with the Clinical Laboratory Standards Institute guidelines (4).

The four colistin susceptible isolates were sub-cultured daily on Mueller Hinton agar (MHA) (Becton, Dickinson and Company, U.S.) containing 1 µg/mL colistin until 40^th^ generation. All generations were tested for colistin MIC’s and alterations in *pmrCAB* and *Lpx* genes. Mutations in *lpxA, lpxC, lpxD* and *pmrCAB* were detected by Sanger Seguencing (Applied Biosystems ABI 3500) using previously described primers (5, 6). ABI files were analyzed in Applied Maths Bionumerics version 7.5 Software (Biomerieux, France). The expressions of *pmrCAB* were studied by qRT PCR (LightCycler 480 II, Roche, Germany) using previously described primers (5, 6). Housekeeping gene was 16SrRNA and *A. baumannii* ATCC 19606 was used for normalization of Ct values. Carbapenamase typing and *mcr-1* positivity was studied by PCR.

The clinical progression of the patient was shown in Figure 1. After initiation of the colistin therapy, two susceptible isolates (K399 and K408) were detected at the 9^th^ and 15^th^ days. The colistin resistant *A. baumannii* (K409) was isolated after 25 days of therapy. All the isolates were found to be clonally related with >95% similarity and positive for *blaOXA-23*. No *mcr-1* was detected. In the colistin resistant isolate (K409), multiple insertions were found in different regions of *pmrA, pmrB, pmrC, lpxA, lpxC,* and *lpxD* genes. The *pmrA, pmrB* and *pmrC* genes were also found to be 1.6, 1.74 and 1.72-fold overexpressed compared to the susceptible K408, respectively.

**Figure 1.**
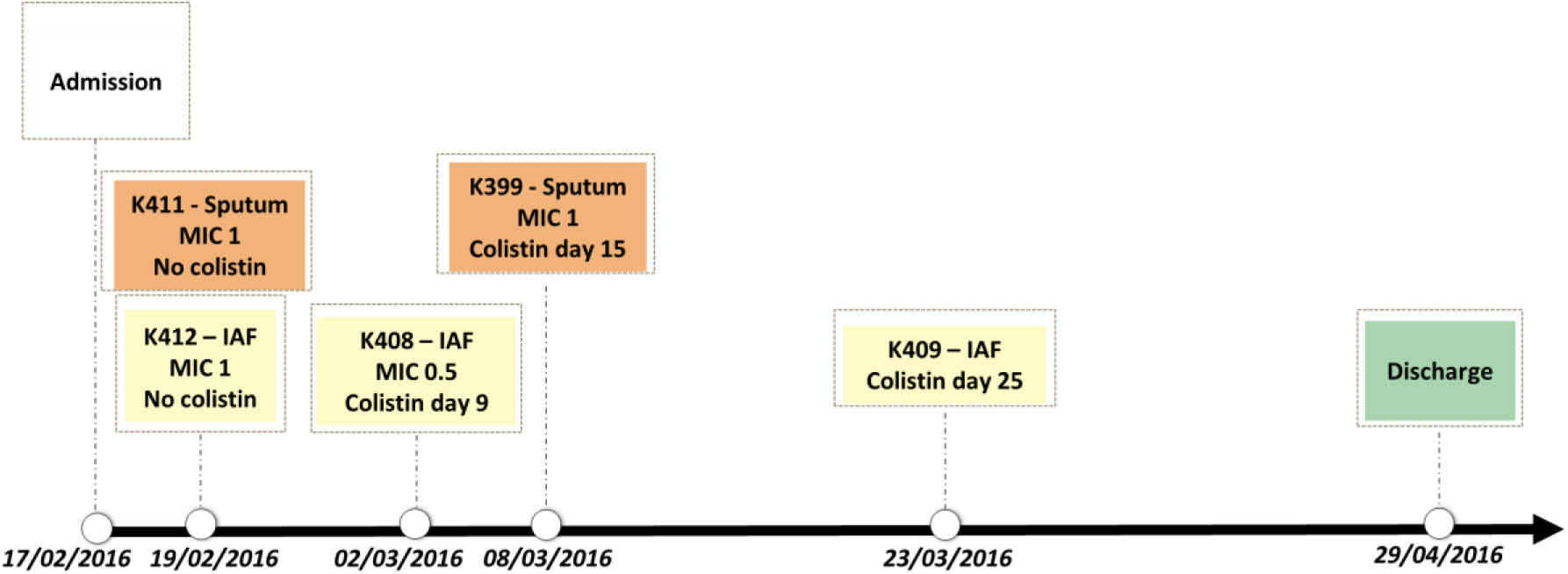
The clinical progression of the patient from admission to discharge of the hospital. The K411 and K412 were isolated before colistin treatment, K408 and K399 were at the 9^th^ and 15^th^ days of the therapy. The colistin resistant K409 was isolated five days after stopping 25 days of colistin therapy.

In the laboratory, the *pmrCAB* expressions of susceptible isolates were found to be significantly induced at the generations that correspond 26^th^ day of colistin exposure which were 11^th^ generation for K399, 17^th^ generation for K408 and 26^th^ day for K412. Colistin MIC’s increased above 2 mg/L after 1^st^ generation and the highest MIC of all generations was 8 mg/L (Figure 2).

**Figure 2:**
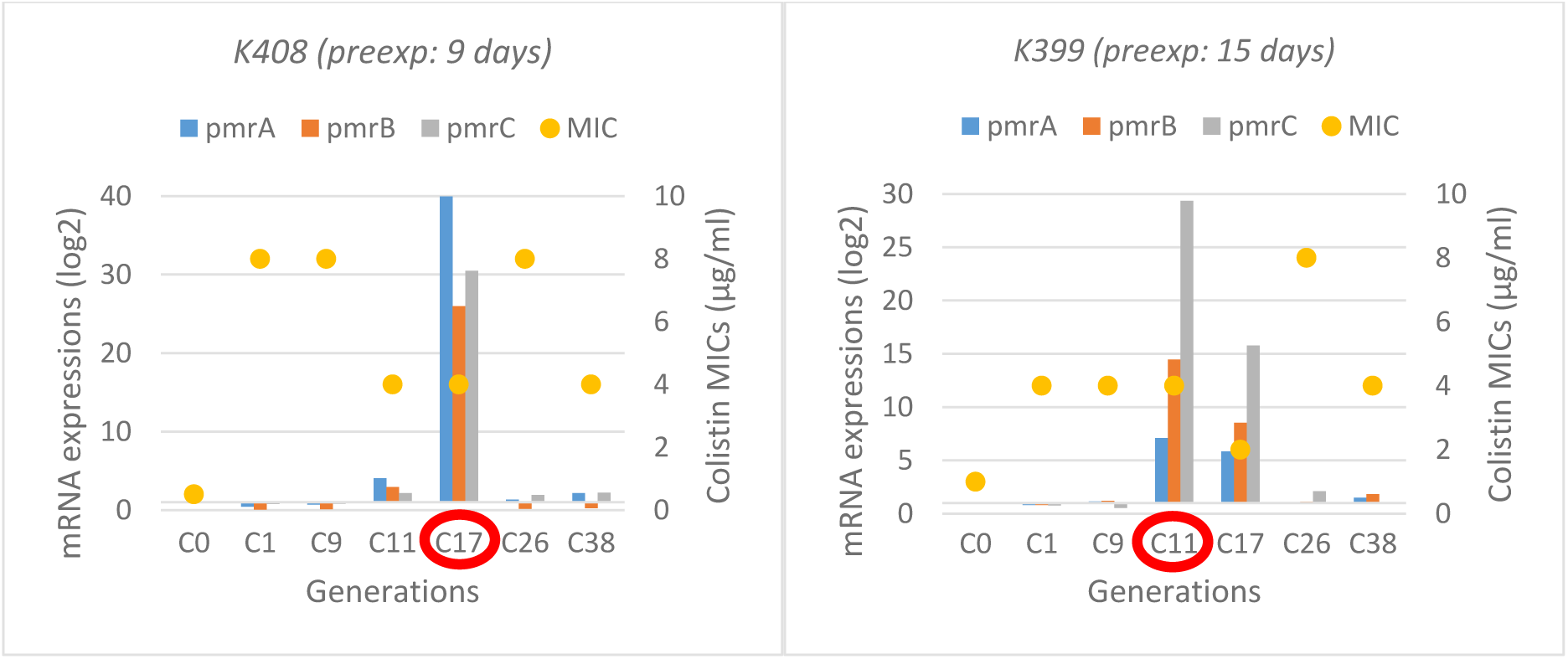
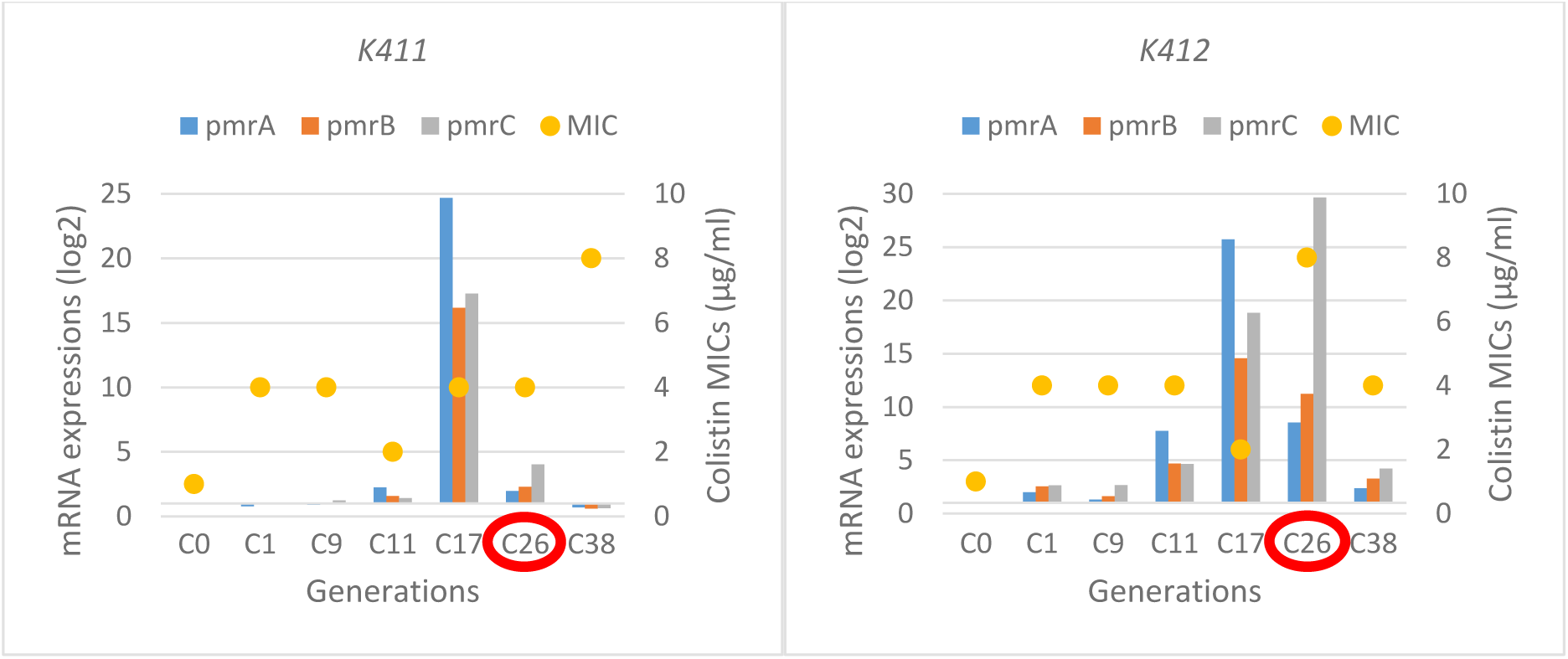
The MIC levels and *pmrCAB* expressions of selected subculture generations of susceptible isolates under colistin exposure. Red circles indicate the generation that corresponds the clinical colistin exposure time

Colistin is one of the last resorts in treatment of MDR *A. baumannii* infections, however, resistance against colistin is increasing rapidly (7). Many researchers reported development of colistin resistance either in laboratory environment or in clinical settings; however, the causes of resistance are still not well known. This study focuses on colistin resistance mechanisms of *A. baumannii* induced by *in-vivo* and *in-vitro* colistin exposure.

In our study, after 25 days of colistin treatment, colistin resistant *A. baumannii* was isolated from intra-abdominal fluid of the patient. Colistin exposure is found to be the main risk factor for development of colistin resistance by previous studies (2, 8). We mimicked the effects of colistin exposure in the laboratory by measuring MIC and expressions of *pmrCAB*. After first passage in the laboratory, colistin MIC increased above the resistance breakpoint (>2 mg/L). (Figure 2). A recent study also reported phenotypically colistin-resistant mutants of *A. baumannii* that was obtained after 24 hours of colistin exposure (9). However, after the 25^th^ day of colistin therapy, we isolated colistin susceptible *A. baumannii* from the patient. Molecular analyses of generations in the laboratory supported this finding and the expressions of *pmrCAB* in generations reached at the highest level at the 11^th^ generation of K399, 17^th^ generation of K408 and 26^th^ generation of K411, which were closely corresponded to the duration of therapy (Figure 2). Development of colistin resistance is an adaptation to environmental stress (10); therefore, it needs a time period for regulation of genetic and metabolic activities. It was shown that mutations and regulation of expression patterns are gained by stress-exposed bacteria in a certain period (11). Lee and colleagues investigated induced colistin resistance in *P. aeruginosa* and found that 6 days of 4 mg/L of colistin exposure was sufficient to gain colistin-resistant phenotype in vitro (12). The expressions of *pmrA, pmrB* and *pmrC* genes of colistin-resistant (K409) isolate were also found to be 1.6, 1.74 and 1.72-fold higher compared to susceptible one, respectively. Beceiro et al reported increased expression of *pmrA* (4- to 13-fold), *pmrB* (2- to 7-fold) and *pmrC* (1- to 3-fold) in resistant *A. baumannii* (6).

We conclude that, duration of colistin exposure is critical for adaptation and establishment of colistin-resistant phenotype. *A. baumannii* can develop a stable colistin-resistant phenotype with elevated MICs and upregulated *pmrCAB* operon after three weeks of colistin exposure.

